# Benchmarking cryo-EM single particle analysis workflow

**DOI:** 10.1101/311324

**Authors:** Laura Y. Kim, William J. Rice, Edward T. Eng, Mykhailo Kopylov, Anchi Cheng, Ashleigh M. Raczkowski, Kelsey D. Jordan, Daija Bobe, Clinton S. Potter, Bridget Carragher

## Abstract

Cryo electron microscopy facilities running multiple instruments and serving users with varying skill levels need a robust and reliable method for benchmarking both the hardware and software components of their single particle analysis workflow. The workflow is complex, with many bottlenecks existing at the specimen preparation, data collection and image analysis steps; the samples and grid preparation can be of unpredictable quality, there are many different protocols for microscope and camera settings, and there is a myriad of software programs for analysis that can depend on dozens of settings chosen by the user. For this reason, we believe it is important to benchmark the entire workflow, using a standard sample and standard operating procedures, on a regular basis. This provides confidence that all aspects of the pipeline are capable of producing maps to high resolution. Here we describe benchmarking procedures using a test sample, rabbit muscle aldolase.

## 1 Introduction

At the Simons Electron Microscopy Center (SEMC) at the New York Structural Biology Center (NYSBC) in New York, NY, our mission is to provide scientific expertise and resources for our users in their studies of biological macromolecules, with a focus on high-resolution structure determination. Our facility is home to seven electron microscopes (EMs), including three 300 kV FEI Titan Krios instruments, all of which are routinely checked for their performance using a series of benchmarking tests. While these checks include standard testing for performance and resolution, typically using a cross grating replica, we believe it is also important to test our systems using a biological sample that scrutinizes the entire workflow from specimen preparation through imaging and image processing. This benchmarking enables us not only to assess any limitations and bottlenecks that might arise, but also allows us to optimize the single particle analysis (SPA) workflow, thus maximizing the throughput and performance of instrumentation and data collection strategies. In addition to the practical advantages of benchmarking, the overall workflow serves as an educational tool for newcomers to cryo electron microscopy (cryo-EM) who wish to learn the SPA workflow using a protein that can be routinely reconstructed to high resolution. Finally, benchmarking tests provide an objective measure to the user that the instrumentation is operating at its top optical efficiency, capable of providing good quality structures, and that any limitations to resolution are thus most likely related to an individual sample.

Benchmarking efforts for SPA are not straightforward for EM labs. This challenge is due in part to the lack of an “industry standard” biological EM specimen and also due to intrinsic variabilities that exist at the specimen purification and grid preparation level. This variability is then coupled with a wide range of data collection and image processing strategies and software choices. An ideal cryo-EM benchmarking standard would be a biological specimen with the following attributes: 1) easily accessible (i.e. commercially available and requiring minimal additional purification), 2) low maintenance sample preparation, 3) biochemically stable over a range of temperatures and time periods, 4) little to no conformational and compositional heterogeneity.

It is also important that the benchmark results in a structure with a sufficiently high resolution, which we consider to be below 3 Å, in order to give confidence in users as to the performance of the instrument, the data collection protocols and the processing pipeline. There are currently 36 unique structures in the EMDataBank at a sub 3 Å resolution that have been obtained by SPA. These include the 465 kD beta-galactosidase at 2.2 Å resolution (Bartesaghi et al. 2015), the 540 kD p97 at 2.3 Å (Banerjee et al. 2016),the 334 kD glutamate dehydrogenase at 1.8 Å (Merk et al. 2016) and the 150 kD aldolase at 2.6 Å (Herzik, Wu, and Lander 2017) as well as larger proteins that have been used as standards in cryo-EM SPA, like the 700 kDa *T. acidophilum* 20S proteasome (Li et al. 2013, Danev, Tegunov, and Baumeister 2017, Campbell et al. 2014, Campbell et al. 2015, Danev and Baumeister 2016) and 440 kDa apoferritin (Russo and Passmore 2014, Rickgauer, Grigorieff, and Denk 2017, Arnold et al. 2017).

In this paper, we present a workflow for single particle reconstruction using a robust and reliable benchmarking standard: rabbit muscle aldolase, a small homotetrameric glycolytic enzyme with a molecular weight of ~150 kDa. Our goal is to present this benchmarking procedure as a step-by-step workflow that can be readily repeated. We show that in order to achieve a sub 3 Å reconstruction of aldolase in a reasonable time frame, ice thickness of 10–20 nm is essential.

## 2 Methods and Materials

### 2.1 Sample Preparation

The sample was prepared as previously described with minor adjustments (Herzik, Wu, and Lander 2017). Briefly, pure aldolase isolated from rabbit muscle (Sigma Aldrich, product #A2714) was solubilized in 20 mM HEPES (pH 7.5), 50 mM NaCl at 3 mg/ml and further purified using a Superose 6 10/300 GL (GE Healthcare) column equilibrated in solubilization buffer. SDS-PAGE analysis was used to confirm sample purity of peak fractions, which were pooled and concentrated to 10 mg/mL and flash frozen in 10 μl aliquots for long term storage. The protein was diluted to 1.5 mg/ml final concentration for grid preparation. Vitrified specimens were prepared by adding 3 μl aldolase (1.5 mg/ml) to freshly plasma cleaned (Gatan Solarus plasma cleaner, 75% argon/25% oxygen atmosphere at 15 Watts for 6 seconds) Au R1.2/1.3 300-mesh (EMS UltrAuFoil®) grids. To minimize the effects of beam induced motion during acquisition, samples were prepared on gold grids (Russo and Passmore 2014). Grids were blotted for 1 s after a 10 s pre-blotting time, then plunge-frozen in liquid ethane using a Leica EM GP instrument (Leica Microsystems), with the chamber maintained at 4°C and 90% humidity.

### 2.2 Microscope Alignment

Complete microscope alignment procedure, based on the FEI on-line manual, were performed during installation using a cross-grating calibration grid (Titan on-line help manual—Alignments, version 2.6 and higher). A minimal subset of the alignments is performed before each daily data collection. These include dark and bright gain corrections and energy filter alignment, performed over vacuum, and beam tilt pivot points and Cs (spherical aberration coefficient) correction, performed at eucentric height and eucentric focus over carbon. Second-order axial coma free alignment and astigmatism minimization was done using the Cs corrector, aligning until A1 (2-fold astigmatism) was less than 10 nm and B2 (coma) was less than 50 nm. Re-tuning of the Cs corrector was performed if the CTF estimation indicated a differential between the major and minor axis of greater than 100 nm. A full tune of the energy filter was carried out daily and energy filter slit was realigned every 60 minutes, managed automatically by Leginon (Suloway et al. 2005). The image distortion after tuning is typically within 0.2 %, and the slit movement was generally +/-1 eV. The goal of these alignments is to verify the presence of Thon rings visible beyond 3 Å resolution in the power spectrum of aligned images collected over amorphous carbon using the same imaging conditions as for the data collection. The eucentric height and eucentric focus are set using Leginon (Suloway et al. 2005) by minimizing movement caused by stage tilt. The beam intensity was kept well within the parallel range of the 3-condenser lens Titan system, with the illuminated beam diameter at least 2–3 times larger than the minimum required for parallel illumination. At a nominal magnification of 130,000x the calibrated beam diameter for parallel illumination is 0.45 – 12.0 μm. In general, the beam diameter was set to be slightly larger than the nominal hole size of 1.2 μm. This helps to ensure that the beam will contact the gold substrate during exposure collection, potentially helping to dissipate charge onto the substrate instead of the sample. Dose rate measurements on the Gatan K2 Summit direct electron detector (DED) were collected to determine whether or not changes to spot size were necessary to achieve the desired dose rate. All high magnification imaging was done in the nanoprobe mode with a 70 μm C2 aperture and a 100 μm objective aperture.

### 2.3 Data Collection

Table 1 summarizes the data collection statistics for three different datasets, 17sep21j, 17nov02c, and 17dec27a. Briefly, data was acquired using a Titan Krios with a spherical aberration corrector and a post-column Gatan Image Filter (GIF) operating in nanoprobe and EF-TEM mode with an extraction voltage of 4250 V, a gun lens setting of 4, a spot size of 6 or 7, a C2 aperture size of 70 μm, an objective aperture size of 100 μm, and an energy filter slit width of 20 eV. The microscope is equipped with a field emission gun operating in the X-FEG module. Data was collected automatically using the *MSI-T2* application in Leginon and all image pre-processing was performed using the Appion pipeline (Lander et al. 2009). Square and sub-square level images were targeted by stage position movement, with a 2 and 5 second pause before imaging, respectively. Drift monitor cutoff was 6 Å/sec. Focusing was performed on the gold substrate, after which four final high-magnification movies were acquired by image shift targeting with a 5 second pause before the first image and 2.5 second pause before each subsequent movie. Final high-magnification movies were taken at a nominal magnification of 130,000x (calibrated pixel size of 0.855 Å at the detector level) and a nominal defocus range of -1.0 to -2.0 μm defocus with the Gatan K2 Summit DED operating in either counting or super-resolution mode. Each movie was acquired over 6000–6600 ms with a frame rate of 5 frames/sec and a dose rate of 8 electrons/pixel/sec. The total cumulative dose for all datasets was in the range of 60–70 electrons/Å^2^.

**Table 1.** Data collection statistics. All datasets were collected on the same Titan Krios equipped with an energy filter and spherical aberration corrector at 300 keV accelerating voltage. Micrographs were collected on the K2 Summit DED, at either counting or super-resolution mode, at a nominal magnification of 130,000x, equivalent to a 0.85 Å pixel size at a dose rate of 8.0 e^-^/pixel/sec. Images were targeted by image shift movement, with a nominal defocus range of -1.0 to -2.0 μm underfocus. The difference in the number of micrographs collected per hour for the 17nov02c and 17dec27a is due to the delayed start of early return on the K2 camera system, which is used to speed up exposure acquisition speeds by outputting only the first few frames of a movie instead of the entire movie so that the camera can continuously collect images.

|  | 17sep21j | 17nov02c | 17dec27a |
| --- | --- | --- | --- |
| **Counting vs Super-resolution** | counting | super-resolution | super-resolution |
| **Exposure time (ms)** | 6600 | 6000 | 6000 |
| **Total dose (e<sup>-</sup>/Å<sup>2</sup>)** | 68 | 63 | 63 |
| **Duration of data collection** | 18 hours | 52 hours | 40 hours |
| **Micrographs collected (per hour)** | ~38 | ~31 | ~40 |
| **# images total** | 699 | 1635 | 1614 |

In addition to our standard data collection workflow, we routinely collect ice thickness measurements for each high magnification movie. This is done by comparing the intensities of images taken without and with the energy filter slit inserted (Rice et al. 2018) (Figure 1D, 1H and 1L).

**Figure 1.**
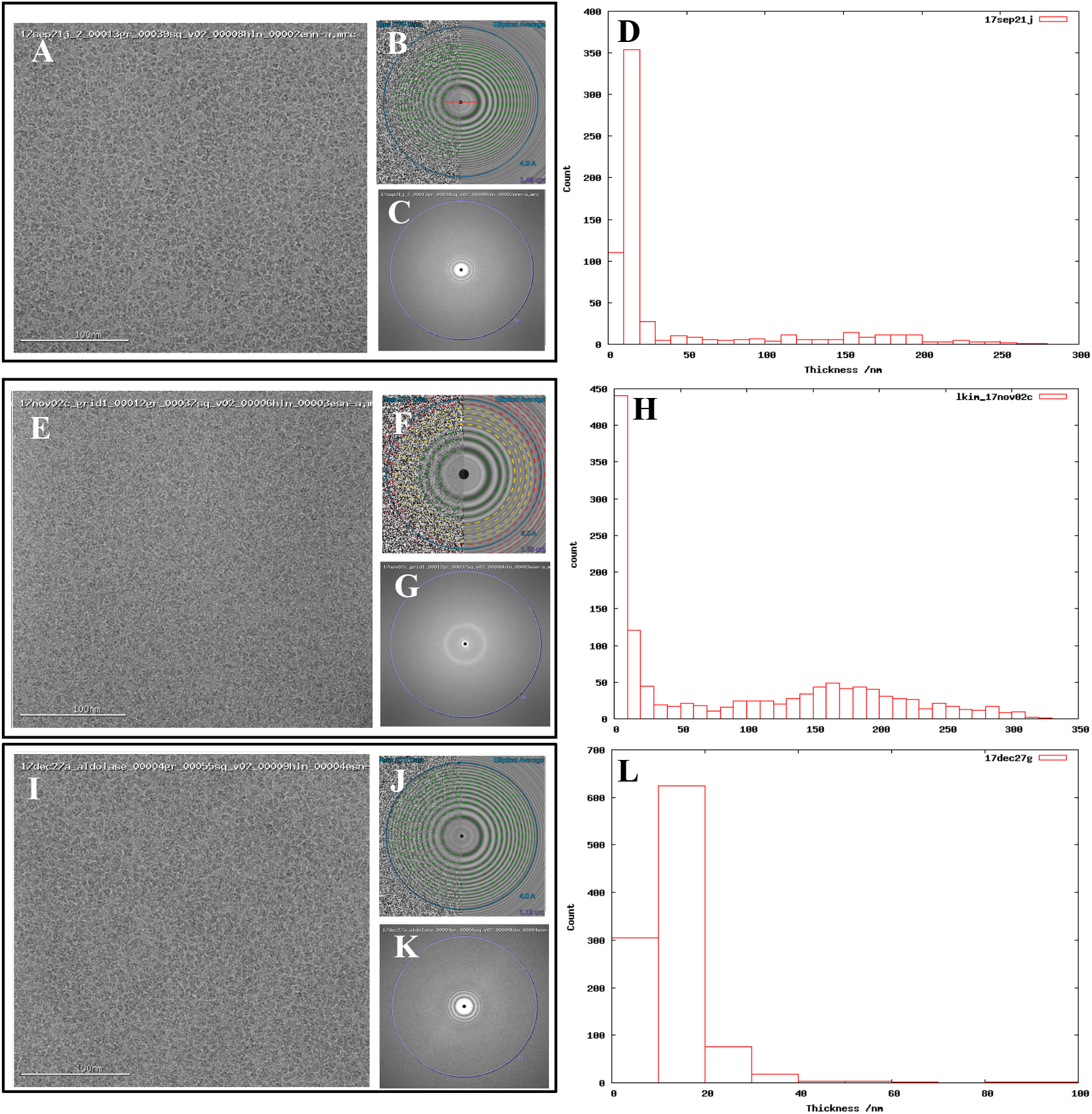
Comparing images from thick versus thin ice. Exemplary images from (a) 17sep21j (#1), (e) 17nov02c (#2), and (i) 17dec27a (#3) datasets. Quantitative metrics such as the estimation of resolution from CTFFindV4 (b, f, & j) and qualitative metrics such as the presence/absence of the water diffraction ring around the 3 Å mark (c, g, & k), and ice thickness measurements of the micrographs (d, h, & 1), should be monitored during data collection. A CTFFindV4 resolution estimation worse than 4 Å and the presence of a strong water diffraction ring are both indicative of thick ice, and areas like this should be avoided. All images were acquired with ~1.5 um defocus. Ice thickness measurements provide a useful metric for data quality (d, h, & 1). Datasets 17sep21j and 17dec27a both contain a majority of images where ice thickness is in the range 0 – 20 nm. The majority of 17nov02c images have thickness in the range 0 – 10 nm thickness (ice that is too thin or completely absent) or very thick ice in the range 100 – 250 nm. The dimensions of aldolase are ~100 Å so this thick ice is more than 20 times more than the longest dimension of the particle.

### 2.4 Concurrent Image Processing

During data collection, images were pre-processed to provide a feedback on image quality. All preprocessing was carried out using the Appion pipeline (Lander et al. 2009). Mechanical and beam-induced motion correction and dose weighting were performed on the raw movies using MotionCor2 (Zheng et al. 2017) using a 5×5 patch size and a B-factor of 100 with 7 iterations. Super-resolution movies were binned by two before frame summation. Whole-image contrast transfer function (CTF) estimation was performed using CTFFind4 (Rohou and Grigorieff 2015). Particle picking was performed within Appion using FindEM template picking (Roseman 2004) with templates generated from images of the same sample acquired on a screening microscope. Box files from the particle picks were generated within Appion and exported for further processing. Subsets were exported for processing during collection and the final full datasets were then processed post-collection.

### 2.5 Post Collection Image Processing

Reference-free 2D classification was performed using cryoSPARC (Punjani et al. 2017) on particles binned by 4 with a box size of 256 pixels. Particles exhibiting secondary structure elements were selected for further processing, including initial model generation, and subsequent 3D classification and 3D auto-refinement using both RELION 2.0 (Kimanius et al. 2016) and cryoSPARC (Punjani et al. 2017). Default processing parameters were generally used. All reported resolutions are based on the 0.143 Fourier shell criterion (Henderson et al. 2012, Scheres and Chen 2012) with all Fourier shell correlation (FSC) curves corrected for the effects of soft-masking by high-resolution noise substitution (Chen et al. 2013). Data processing statistics, including number of particles and average processing times, are described in Table 2.

**Table 2.** Processing and Reconstruction statistics. Datasets were sorted based on three methods, using all micrographs, using micrographs with < 25 nm ice thickness and using only the first few hundred micrographs. This is to test whether it is more important to collect and process data based on the quantity of data (all micrographs), quality of data (< 25 nm ice thickness), or time spent on data collection and processing (first few hundred micrographs). All 2D and 3D processing was performed using Cryosparc.

|  | 17sep21j | 17sep21j | 17sep21j | 17nov02c | 17nov02c | 17nov02c | 17dec27a | 17dec27a | 17dec27a |
| --- | --- | --- | --- | --- | --- | --- | --- | --- | --- |
| **Sorting method** | All img | < 25 nm ice thickness | 1<sup>st</sup> 500 img | All img | < 25 nm ice thickness | 1<sup>st</sup> 700 img | All img | < 25 nm ice thickness | 1<sup>st</sup> 382 img |
| **Dataset number** | #1 | #1a | #1b | #2 | #2a | #2b | #3 | #3a | #3b |
| **Duration of data collection** | 18.0 hours | 18.0 hours | 13.5 hours | 52.0 hours | 52.0 hours | 24.0 hours | 40.0 hours | 40.0 hours | 10.0 hours |
| **# images total** | 699 | 699 | 699 | 1635 | 1635 | 1635 | 1614 | 1614 | 1614 |
| **# images used** | 699 | 535 | 500 | 1635 | 63 | 700 | 1614 | 1108 | 382 |
| **# picks** | 642K | 491K | 256K | 1,380K | 60K | 685K | 1,214K | 975K | 234K |
| **Duration of 2D classification** | 0.8 hours | 0.4 hours | 0.3 hours | 1.3 hours | 0.3 hours | 0.4 hours | 1.1 hours | 1.1 hours | 0.4 hours |
| **# particles after 2D** | 373K | 374K | 133K | 464K | 26K | 198K | 498,000 | 369K | 87K |
| **% particles after 2D** | 58% | 76% | 52% | 34% | 43% | 29% | 41% | 38% | 37% |
| **Duration of 3D classification** | 1.2 hours | 1.7 hours | 0.9 hours | 3.9 hours | 0.5 hours | 0.5 hours | 6.3 hours | 5.1 hours | 0.8 hours |
| **# particles into refinement** | 219K | 124K | 62K | 204K | 22K | 75K | 205K | 187K | 87K |
| **% particles into refinement** | 59% | 33% | 47% | 44% | 85% | 38% | 41% | 52% | 100% |
| **Duration of refinement** | 0.9 hours | 0.9 hours | 0.9 hours | 1.9 hours | 0.5 hours | 0.5 hours | 1.5 hours | 1.4 hours | 0.7 hours |
| **Ice thickness range** | 10 – 20 nm | 10 – 20 nm | 10 – 20 nm | 100 – 250 nm | 100 – 250 nm | 100 – 250 nm | 10 – 20 nm | 10 – 20 nm | 10 – 20 nm |
| **Total processing time** | 2.9 hours | 3.0 hours | 2.0 hours | 7.1 hours | 1.2 hours | 1.4 hours | 8.9 hours | 7.6 hours | 1.9 hours |
| **Resolution (global)** | 2.5 Å | 2.5 Å | 2.8 Å | 3.0 Å | 3.5 Å | 4.6 Å | 2.4 Å | 2.4 Å | 2.8 Å |

Figure 1 shows exemplary images from the datasets #1, #2, and #3. Figure 2 shows processing results from dataset #1 and #2, including 2D class averages, Euler plots, FSC curves, 3D maps, and ice thickness plots.

**Figure 2.**
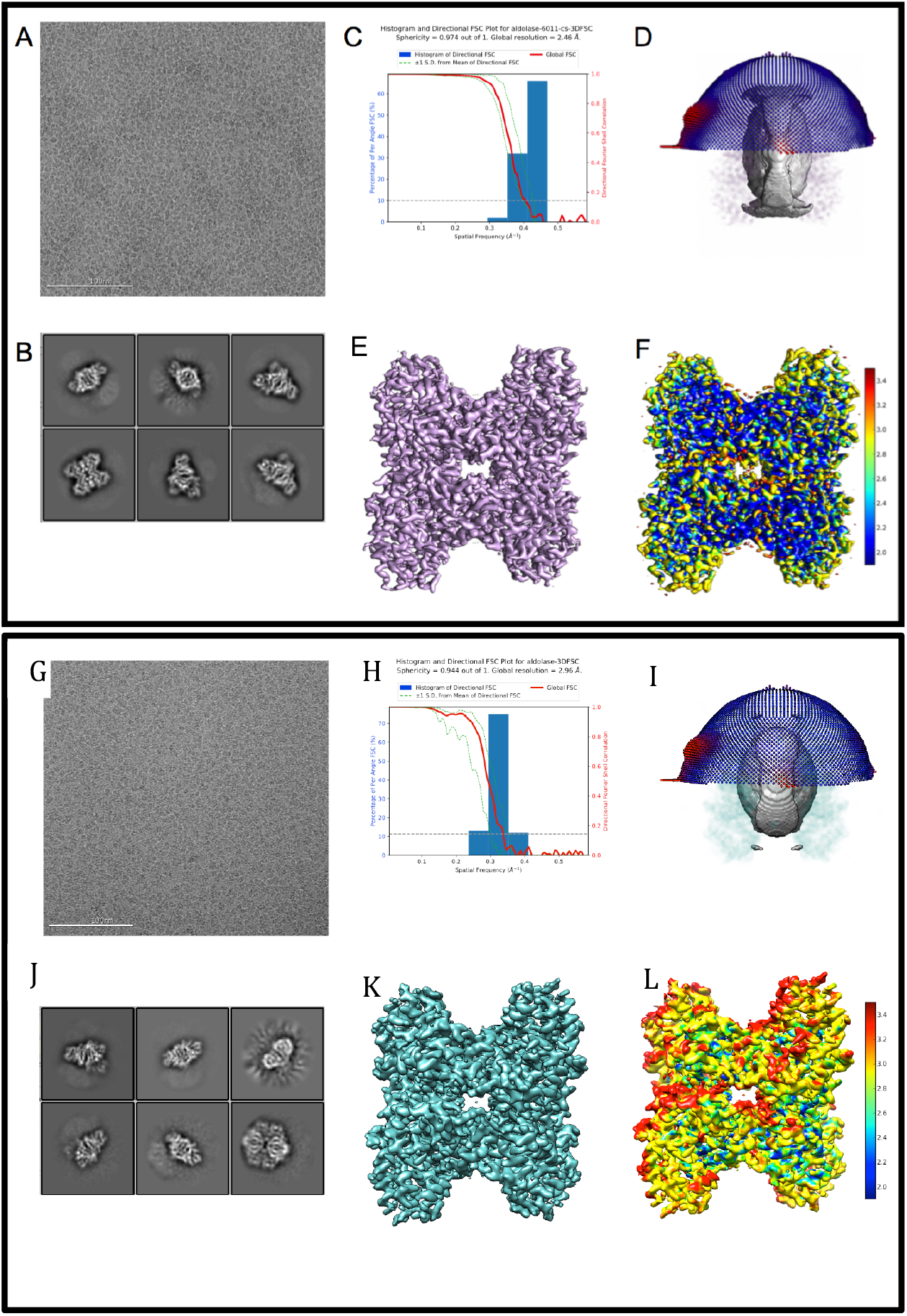
Comparing 3D reconstructions from thick versus thin ice. 2D and 3D processing results from 17sep21j (dataset #1) and 17nov02c (dataset #2) which yielded maps at 2.5 Å and 3.0 Å resolution, respectively. Dataset # 1 has thinner ice in the raw micrographs, ranging from 10 – 20 nm thick, whereas dataset #2 has thicker ice, ranging from 100 – 250 nm thick. A raw micrograph (a & g), 2D classification (b & h), FSC plot (c & i), sphericity (d & j), 3D map (e & k) and local resolution map (f & 1). Both datasets have about 200,000 particles contributing to the final refinement but dataset #1 is both qualitatively and quantitatively better than dataset #2.

## 3 Results and Discussion

We_describe three unique data sets that were processed based on three different sorting criteria: the first few hundred images collected for the session, all images collected in the session, images sorted by ice thickness < 25 nm, for a total of nine experiments (Table 2). The details of data acquisition and processing are provided in Table 1 and 2, respectively.

While all three data sets result in 3 Å or better maps, the major variable in terms of data and map quality was ice thickness. For grids with very thin ice, sub 3 Å maps can be obtained in a much shorter time period and with fewer images than from grids with much larger ice thickness. Use of image shift navigation as opposed to stage position navigation is our preferred mode of data collection as it helps to maximize acquisition throughput, and moderate amounts of image shift do not affect results at the targeted resolution (Cheng et al. 2018).

From experience (and personal communication with the Lander lab) we have concluded that it is very important to maximize the number of particles packed into each hole, while avoiding particle overlap and aggregation. This close packing provides a more accurate CTF estimation of each image since the protein contributes a high signal to the power spectrum of the image. We also hypothesize that the densely packed protein is instrumental in achieving a very thin ice layer, as it may help to retain a thin layer of liquid across the hole. Dense packing of the protein is concentration dependent and can lead to multiple layers of particles (Noble et al. 2017). This is refractory to high-resolution goals as two layers of protein result in a whole-image defocus estimation averaged between the two layers, thus limiting the resolution of each particle depending on their distance from the midway point.

Table 2 shows the results of processing the 17sep21j dataset (#1, #1a and #1b) in three different ways including all 699 images (dataset #1), sorting based on including only images with < 25 nm ice thickness (dataset #1a) and using only the first 500 images (dataset #1b). We found that resolution of 2.5 Å can be achieved either by using all images or by using only those from the thinnest ice. Micrographs coming from the thinnest ice yield a higher resolution final reconstruction and thus limiting image acquisition to areas of thin ice is clearly a more efficient strategy than brute force processing of the largest number of images. The 17sep21j dataset is near-perfect in that it can yield a sub 3 Å reconstruction in under 24 hours, regardless of how the data is sorted because of the majority of images coming from a very thin ice.

The 17nov02c (#2, #2a and #2b) dataset is representative of the type of data collection that should be avoided if possible. This dataset required a large block of microscope time (52 hours), processing time and computational resources (over 1.3M particles before 2D classification). While dataset #2 yielded a 3.0 Å map it required a total data collection and processing duration of ~60 hours. Ideally, a benchmarking test should be accomplished in less than 24 hours. Also, the sorted data from this dataset (#2a and #2b) both provided reconstructions worse than 3.0 Å, due to the very small number of particles that are in thin ice.

The 17dec27a (#3) dataset yielded a similarly high-resolution data as the 17sep21j (#1) dataset, but it required more than twice the length of microscope and processing time, 49 hours versus 21 hours. Both datasets contributed about 200K particles to the final refinement, but 17dec27a (#3) started with more than twice the number of micrographs compared to 17sep21j (#1) (1614 versus 699 micrographs).

Table 3 ranks the datasets by nominal resolution. We find that all sub 3 Å reconstructions are derived from datasets with an ice thickness range of 10–20 nm, independent of how the data was sorted indicating that ice thickness is a primary driver of data quality for the SPA of aldolase. Quantitative metrics like ice thickness measurements, or qualitative metrics such as the presence of an ice ring in the power spectrum of the image should be used to guide data collection strategy during collection. The 17sep21j and 17dec27a datasets had ice thickness measurements on average around 10–20 nm, whereas the 17nov21j ranged from 100-250 nm. Rabbit muscle aldolase, a 150 kDa homotetramer, has unit cell dimensions of 82.8 x 100.6 x 84.5 Å, so that individual particles are readily visible in ice 10–25 nm thick, but once embedded in ice that is 100–250 nm thick, contrast is much worse and individual particles are difficult to identify. In addition to the loss of contrast in the images, once the ice thickness is nearly 10 times that of the longest length of the particle of interest, it is likely that multiple layers of particles are present in the image, adhering to either side of the exposed air-water interface (Noble et al. 2017) which would also interfere with the possibility of getting a high resolution reconstruction.

**Table 3.** Ranking of datasets based on data collection and processing time and resolution. Datasets were ranked primarily on total data collection + processing time and secondarily on nominal resolution. Six out of nine datasets went to < 3 Å. We find that all < 3 Å reconstructions come from datasets with 10 – 20 nm ice thickness and that more than half of those < 3 Å datasets were acquired in under 24 hours. Datasets that do not go < 3 Å had ice thickness measurements ranging from 100 – 250 nm.

| Ranking | Dataset | Total collection + processing time | resolution | # particles | Ice thickness range |
| --- | --- | --- | --- | --- | --- |
| 1 | #3b | 11.9 hours | 2.8 $\text{\AA}$ | 87K | 10 – 20 nm |
| 2 | #1b | 15.5 hours | 2.8 $\text{\AA}$ | 62k | 10 – 20 nm |
| 3 | #1 | 20.9 hours | 2.5 $\text{\AA}$ | 219k | 10 – 20 nm |
| 4 | #1a | 21.0 hours | 2.5 $\text{\AA}$ | 124k | 10 – 20 nm |
| 5 | #3a | 47.6 hours | 2.4 $\text{\AA}$ | 186k | 10 – 20 nm |
| 6 | #3 | 48.9 hours | 2.4 $\text{\AA}$ | 205k | 10 – 20 nm |
| 7 | #2 | 59.1 hours | 3.0 $\text{\AA}$ | 204k | 100 – 250 nm |
| 8 | #2a | 53.2 hours | 3.5 $\text{\AA}$ | 22k | 100 – 250 nm |
| 9 | #2b | 25.4 hours | 4.6 $\text{\AA}$ | 75K | 100 – 250 nm |

We conclude that the most important factor in reaching a sub 3 Å map in less than 24 hours for our aldolase benchmark specimen is to have grids with ice thickness in the range of 10–20 nm. Use of a Cs-corrected system is not required for achieving these results as we have been able to replicate similar results using a well-aligned, non Cs-corrected systems. Similar results were achieved from the super-resolution and counting mode datasets, implying that it is not necessary to collect data in super-resolution mode to produce a sub 3 Å reconstruction. We also note that on-the-fly data processing, which includes frame alignment, CTF estimation, and particle picking, is critical to maximizing the quantity and quality of data collected. Real-time feedback of the data helps guide the data collection strategy, allowing the user to be more critical about which regions of a grid, square, and hole to collect in (based on information on particle density, distribution, ice thickness, etc.), how long to pause between images (based on the motion correction plots), and how much defocus to apply (depending on how much contrast is visible in the aligned movies).

In summary, we show that with a commercially available protein and minimal biochemical purification, it is possible to prepare grids for characterizing microscopes at high resolution. While our protocol was tested on a Titan Krios microscope equipped with a K2 detector, this protocol could easily be adapted to other workflows (e.g. EPU or SerialEM) and microscope/detector combinations. Having a set of standard samples used by many EM labs will be generally useful for the field.

## Author Contributions

LK, Performed sample preparation, microscope alignment, and data collection;; Wrote the manuscript.

WR, Performed ice thickness measurements and analysis; Edited the manuscript.

EE, Performed data collection & image processing; Edited the manuscript.

MK, Performed microscope alignment and data collection; Edited the manuscript.

AC, Edited the manuscript.

AR, KJ, & DB, Performed sample preparation and data collection.

CP, Conceived and designed experiments; Edited the manuscript.

BC, Conceived and designed experiments; Edited the manuscript.

## Acknowledgments

The authors wish to thank Dr. Gabriel Lander (Scripps Research Institute) for helpful insight and discussions.

## Data Availability Statement

The datasets #1, 1a, 1b, 2, 2a, 2b, 3, 3a, &3b for this study can be found in the Electron Microscopy Data Bank (EMDB) in the form of EM maps. Their accession codes are:

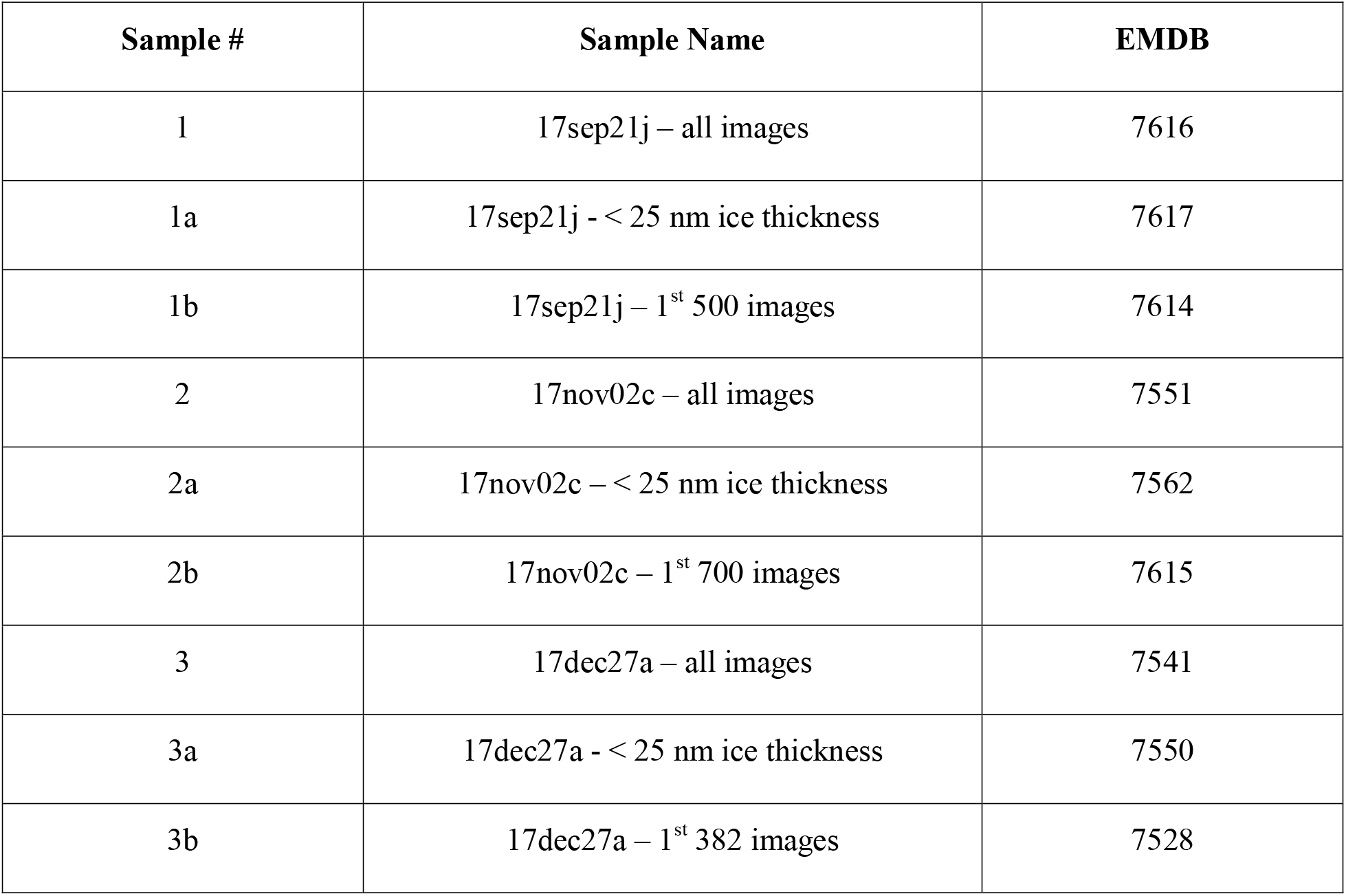

| **Sample #** | **Sample Name** | **EMDB** |
| --- | --- | --- |
| 1 | 17sep21j – all images | 7616 |
| 1a | 17sep21j - < 25 nm ice thickness | 7617 |
| 1b | 17sep21j – 1<sup>st</sup> 500 images | 7614 |
| 2 | 17nov02c – all images | 7551 |
| 2a | 17nov02c – < 25 nm ice thickness | 7562 |
| 2b | 17nov02c – 1<sup>st</sup> 700 images | 7615 |
| 3 | 17dec27a – all images | 7541 |
| 3a | 17dec27a - < 25 nm ice thickness | 7550 |
| 3b | 17dec27a – 1<sup>st</sup> 382 images | 7528 |

